# Context-dependent odor learning requires the anterior olfactory nucleus

**DOI:** 10.1101/2020.01.06.895961

**Authors:** Max Levinson, Jacob P. Kolenda, Gabriella J. Alexandrou, Olga Escanilla, David M. Smith, Thomas A. Cleland, Christiane Linster

**Author notes:** These authors contributed equally to this work (Max Levinson, Jacob P. Kolenda). The authors declare no competing financial interests. The authors are thankful to Matthew Lewis and Norma Hernandez for discussions and assistance with experiments.

## Abstract

Learning to associate the context in which a stimulus occurs is an important aspect of animal learning. We propose that the association of an olfactory stimulus with its multisensory context is mediated by projections from ventral hippocampal networks (vHC) to the anterior olfactory nucleus (AON). Using a contextually-cued olfactory discrimination task, rats were trained to associate two olfactory stimuli with different responses depending on visuo-spatial context. Temporary lesions of the AON or vHC impaired performance on this task. In contrast, such lesions did not impair performance on a non-contextual olfactory discrimination task. Moreover, vHC lesions also impaired performance on an analogous contextually-cued texture discrimination task, whereas AON lesions affected only olfactory contextual associations. We describe a distinct role for the AON in olfactory processing, and conclude that early olfactory networks such as the olfactory bulb and AON function as multimodal integration networks rather than processing olfactory signals exclusively.

**Significance statement:** Contextual information has long been known to play a key role in cognitive functions such as memory and decision making. We here show the contextual modulation of neural information in early primary sensory networks and its effects on contextually conditional learned behavior. We propose that projections from ventral hippocampus to anterior olfactory nucleus convey contextual information to the early olfactory system, modulating sensory representations and olfactory perception. Using behavioral pharmacology and computational modeling, we show how established network structures can mediate multimodal information and use context to make olfactory decisions.

## Introduction

Learning to associate the context in which a stimulus occurs is an important aspect of animal learning. We here show that, in the olfactory system, learning to associate a given odor stimulus with different behavioral responses depending on visuospatial context requires hippocampal as well as anterior olfactory nucleus (AON) function. We describe a novel role for the AON in olfactory processing, and conclude that early olfactory networks such as the olfactory bulb and AON operate as multimodal integration networks rather than processing olfactory signals exclusively.

The AON, a two-layered cortical structure located between the olfactory bulb (OB) and the piriform cortex (PC), is extensively interconnected with related brain regions such as the OB, PC, and hippocampus, among others (Aqrabawi & Kim 2018b, Brunjes et al 2005, Haberly & Price 1978, van Groen & Wyss 1990). It is well established that the AON receives odor stimulus-gated information from OB, PC, and other olfactory structures (Brunjes et al 2005). Neurons in the AON *pars externa* exhibit a consistent spatial patterning of odor responses that is reproducible between individuals, comparably to OB principal neurons and unlike PC pyramidal cells. In contrast, in the AON *pars principalis*, odor-activated neurons are widely distributed without clear spatial patterning, comparable to responses in PC, and respond robustly only to higher odorant concentrations (Kay et al 2011). However, despite its prominent location within the olfactory network, the functional roles of the AON in olfactory processing remain largely unclear.

Recently, work by Oettl et al. (2016) has described a prominent role for the AON in social odor recognition via feedback projections from the AON to the OB. Briefly, in social situations, the activation of abundant oxytocin receptors activates AON pyramidal cells, which then modulate OB processing via their projections onto OB interneurons. This modulation by AON regulates the duration, and possibly the specificity, of a social odor memory (Linster & Kelsch 2019). Aqrabawi et al. (2016) also reported a contribution to odor processing by AON projections to the OB. By temporarily activating or inactivating pyramidal cells in the AON, they showed that AON activity influences odor detection only at low concentrations; detection of weak odors was inhibited by AON activation and enhanced by AON inactivation. In a follow-up study, the same group showed that hippocampal activation of the AON network is required for mice to associate odors and spatial contexts in odor-object memory paradigm in which mice investigated a known odor in a known location less than in a novel location. In contrast, mice performed similarly to controls when a simple odor object recognition task without spatial information was employed (Aqrabawi & Kim 2018a). At the neurophysiological level, Rothermel and Wachowiak (2014) showed that feedback projections from AON to OB are modulated by behavioral state and by the activation of neuromodulatory nuclei, suggesting a multifunctional aspect to this modulation.

In addition to receiving inputs from multiple olfactory networks, hippocampal areas CA1 and the subiculum project ipsilaterally to the AON (Aqrabawi & Kim 2018b, Brunjes et al 2005, Haberly & Price 1978, van Groen & Wyss 1990), such that the ventralmost region of the hippocampus innervates mostly the medial AON, with progressively more dorsal regions innervating more lateral locations within the AON. Experimental activation of fibers projecting from ventral hippocampus (vHC) area CA1 to the AON replicated behavioral effects evoked by activation of the AON itself (Aqrabawi et al 2016), suggesting that this connection is primarily excitatory. The hippocampus is known to be involved in representing spatial context, typically defined as the set of continuously present background cues that comprise the spatial environment. Notably, whereas the hippocampus is not required for many classical and operant conditioning tasks, it becomes critical when a temporal delay or a contextual memory component is introduced (e.g., (Butterly et al 2012, Kim et al 1992, Sill & Smith 2012). The latter observations suggest that hippocampal firing patterns represent aspects of the spatial context, such that the export of these patterns to other brain regions could prime memories that are associated with that context, including olfactory memories (Bulkin et al 2016, Smith & Bulkin 2014). Indeed, odor-context associations, as exhibited by mice searching for a missing odor in a familiar environment, were impaired when feedback inputs to the OB, or projections from the vHC to the AON, were pharmacologically inhibited (Aqrabawi & Kim 2018a, Mandairon et al 2014). These data strongly suggest that contextual information is communicated to the olfactory system via the AON, and that projections from vHC to AON contribute to this communication. For example, rats can readily learn to perform olfactory discrimination tasks in a context-dependent manner (i.e. A+/B-in one context and A-/B+ in a different context), whereas rats with vHC lesions are severely impaired in this task (Komorowski et al 2013). Given the prominent projection from vHC to AON, the AON is strategically positioned to communicate contextual information to early olfactory sensory networks such as the OB.

We here show that a functioning AON is necessary for rats to express a previously learned contextually-cued odor discrimination task. In contrast, neither non-contextually cued odor discriminations nor contextually cued texture discriminations required a functioning AON. We then show that multiple separate generalization gradients around the same odorants can co-exist, and be selectively behaviorally expressed according to visuospatial context, if and only if AON function is unimpaired. Based on the hypothesis that generalization among odorants is mediated by OB odor representations, we use a large-scale computational model of OB and AON to test to what extent the established anatomical circuits of this network could mediate these behavioral effects. The model indicates that AON inputs to the OB can modulate bulbar odor representations in a contextually-dependent manner that is predictive of these behavioral results.

In summary, we propose a novel role for AON in conveying behaviorally relevant contextual information to the early olfactory system.

## Methods

### Behavioral experiments

#### Subjects and Surgeries

Nine male Long-Evans rats, purchased from Charles River Laboratories (Wilmington, MA) and weighing between 400 and 530 grams before training, were used for behavioral experiments. One cohort of six rats was used for Experiments 1a, 1b and 2; four of these six were used for Experiment 1c. A second cohort of three rats was used for Experiment 3. Water was freely provided in the home cages at all times. Rats were kept on a reversed 12:12 light-dark schedule, and all behavioral training and testing took place under dim red light during their dark cycle. During behavioral training and testing, rats were food-restricted to 80 – 85% of their free-feeding weights. Rats were pair-housed up until surgery, after which they were separated into individual cages. Guide cannulae for drug infusions were surgically positioned bilaterally in the AON and vHC as follows: AON: 5.2 mm rostral and ±1.5 mm lateral to bregma, 2.4 mm ventral to the cortical surface; vHC: −5.3 rostral and ±5.0 mm lateral to bregma, 6.6 mm ventral to the cortical surface. Cannula placement was confirmed in a subset of brains after completion of the experiment.

#### Drug infusions

Saline (0.25 uL at 0.1 uL/min) or muscimol (0.25 uL of 0.5 mg/mL at 0.1 uL/min) were bilaterally infused into the AON or vHC using infusion needles 1 mm longer than the implanted guide cannulae. The infusion needle was left in place for 10 minutes after the end of the infusion, and behavioral experiments began 30 minutes after the end of the infusion.

#### Odors

Two different odor sets, each comprised of a pair of household spices (cumin and oregano, mustard and anise seed), were used to train the rats on the contextually-cued conditional discrimination task (Experiment 1a), the non-contextually cued control task (Experiment 1b) and the simultaneous odor/texture task (Experiment 3). A series of straight chain aliphatic aldehydes ranging from 3 to 8 carbons (Cleland et al 2002, Cleland et al 2009) was used for non-contextual generalization testing (Experiment 2a), and a series of straight chain aliphatic butyrates ranging from 2 to 7 carbons was used for contextually modulated generalization testing (Experiment 2b). These odorants were diluted in mineral oil to approximate vapor-phase partial pressures of 1.0 Pa (odor dilutions are shown in Table 1). Odor stimuli were presented to rats in two ceramic ramekins filled with scented bedding in which sucrose reward pellets were buried. Household spices were mixed into the bedding ahead of time, whereas aldehydes and butyrates were pipetted (60 uL) into the bedding prior to the trial.

**Table 1.** Odor sets and vol/vol dilutions used to generate theoretical vapor-phase partial pressures of 1.0 Pa for Experiments 2a and 2b.

| **Experiment 2a** |  |  |  |  |  |  |
| --- | --- | --- | --- | --- | --- | --- |
| Odors | Propanal (C3) | Butanal (C4) | Pentanal (C5) | Hexanal (C6) | Heptanal (C7) | Octanal (C8) |
| %vol.vol | 0.0004 | 0.002 | 0.007 | 0.02 | 0.15 | 0.22 |
| **Experiment 2b** |  |  |  |  |  |  |
| Odors | Ethyl (C2) butyrate | Propyl (C3) butyrate | Butyl (C4) butyrate | Pentyl (C5) butyrate | Hexyl (C6) butyrate | Heptyl (C7) butyrate |
| %vol.vol | 0.002 | 0.05 | 0.17 | 0.57 | 1.63 | 4.57 |

#### Behavioral Shaping

Rats were first trained on a digging task developed for olfactory studies by Linster and Hasselmo (1999). In this task, sugar rewards were buried in small ramekins filled with bedding. Initial dig training took place in a black Plexiglas chamber separated into two compartments by a removable divider. Each training session lasted for 20 trials. For each trial, two dishes were presented in randomized locations, each scented with one of the two odorants, and the rat was given the chance to dig in one. Scored digging behavior consisted of displacement of bedding by either the nose or feet. Upon retrieval of the reward (in correct trials) or cessation of digging (in incorrect trials) the dishes were removed and the rat returned to the waiting area. The reward then was replenished if necessary, the dishes replaced in newly randomized locations, and the next trial began. Shaping was completed once an animal retrieved the sugar reward buried in the bedding in at least 18 out of 20 trials for three successive days. Odorants used for dig training were different from the odorants used for the experimental studies described.

#### Contextually-cued conditional odor discrimination task training

Task training involved a procedure similar to that used for contextually-cued olfactory discrimination in Komorowski et al (2013). A wooden chamber was separated into three compartments: a *white* side, a *black* side, and a central waiting area. The entirety of the white side was painted white and the walls were wiped down with scented baby wipes, while the black compartment was painted black and wiped with unscented baby wipes. In this way the two contexts were distinguishable by visual hue and ambient odor. The task consisted of learning a conditionally-cued discrimination rule using spatial context (i.e., the *black* or *white* side of the box) as the cue and the two odorants as stimuli, such that a dish scented with a certain odor was rewarded in one context but not in the other. After completing dig training, each rat was trained to perform the conditionally-cued discrimination task twice, once with each pair of household spices, with the order of these odorant sets counterbalanced among rats. The pairing of the context and the rewarded odor within each set was also counterbalanced. Rats were trained daily for one session of 40 trials until attaining a criterion of 80% correct trials for two days in a row. Training proceeded according to the following schedule: 40 trials per day on each side until criterion, followed by 20 trials on each side per day, then 10 trials alternating on each side, then 5 trials alternating, until the final stage during which the 40 trials were randomized between sides.

#### Experiment 1: Role of the AON in contextually-cued conditional odor discrimination

To test whether AON inactivation impaired the expression of a learned contextually-cued conditional odor discrimination, rats were bilaterally infused with saline or muscimol into the AONs 30 minutes prior to being tested on 40 randomized trials in the *black* and *white* contexts (Experiment 1a; Figure 1A_i_). Subsequently, as a control, the same rats were trained on a simple odor discrimination task in the same box used for shaping (Figure 1A_ii_), without reference to visuospatial context, and were then infused with muscimol or saline into the AONs and tested (Experiment 1b). To measure the extent to which AON activity is specific to contextually-cued *olfactory* discrimination tasks, a subset of four rats then were trained on a non-olfactory contextually-cued conditional discrimination task. Specifically, instead of using odorants as the discriminative cue, rats were trained to dig in two different types of bedding (corn bedding and paper bedding), with the reward being buried in the paper bedding on one side of the box and in the corn bedding on the other side. Once the rats had reached training criterion, they were infused bilaterally with muscimol or saline into the AONs and then tested on their contextual texture discrimination performance (Experiment 1c). Rats were each run twice under each drug condition; the order of drug infusions was counterbalanced among rats.

**Figure 1.**
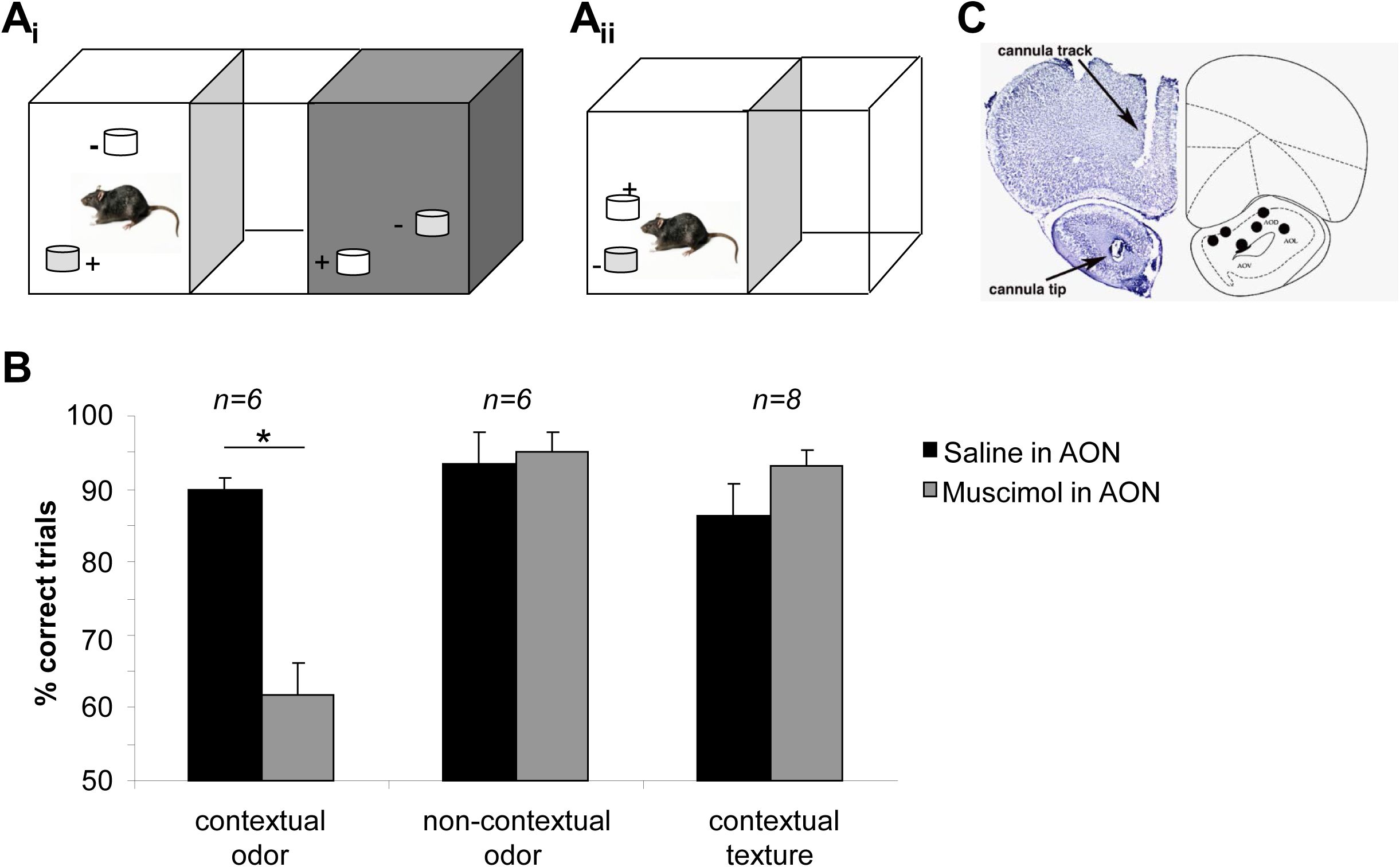
An intact AON is required to express a well learned contextually-cued olfactory discrimination task. **Ai**. Rats were trained in a three-chambered wooden box with a *black*, a *white* and a central *start* chamber of natural wood color. The valence of paired odorants or textures was reversed in the two sides of the box. **Aii**. Rats were trained on a non-contextually cued odor discrimination in a Plexiglas box with a *start* chamber and a *test* chamber. Odor valence was the same over the course of training and testing for a given rat. **B**. The graph shows the percentage (+/-standard error) of correct trials during test days after infusion of saline or muscimol into the AONs. Rats were tested in the contextually-cued odor discrimination task (contextual odor), an odor discrimination task without contextual cues (non-contextual odor) and a texture discrimination task with contextual cues (contextual odors). * depicts a significant difference between drug conditions for a given experiment. **C.** Histologically verified positions of cannula tips in the AON.

#### Experiment 2: Role of AON in perceptual generalization between odorants

To test whether temporary lesions of the AON affect odor perception *per se* (Experiment 2a), six rats were trained to discriminate between pentanal (C5) and hexanal (C6) in the same apparatus used for dig training (Figure 1Aii; non contextually-cued task). For each rat, one odor was rewarded and not the other, with the rewarded odor counterbalanced between rats. Training took place over two sessions of 20 trials with odors C5 and C6, as the rats quickly reached criterion in this non-contextual task. After reaching criterion, on a given testing day, 20 training trials were followed by bilateral saline or muscimol infusions into the AONs. Following a 30 minute resting period to allow the drug to diffuse throughout the AONs, rats were placed back into the waiting area and then given 3 different test trials per day. During test trials, one dish contained unscented bedding, one dish contained bedding scented with any one of the odors in the series C3 – C8 (the *test odor*), and neither dish was rewarded. The orders of test trials were randomized among rats. Rats were given 30 seconds to approach and were allowed to dig in both dishes. The time spent digging in the test dish, a measure for generalization between a previously rewarded odor and a novel odor (Cleland et al 2002, Cleland et al 2009, Linster & Hasselmo 1999, Linster et al 2002), was recorded using a stopwatch.

We then tested the same rats on generalization between odorants after being trained on a contextually cued task using butyl and heptyl butyrate as odors (Figure 1Ai; Experiment 2b). Rats were first trained on the contextually-cued conditional odor discrimination task using the odor pair C4 (butyl butyrate) and C5 (pentyl butyrate). The context/valence pairings were pseudorandomized across subjects, such that for one rat C4 would be rewarded on the white side and C5 in the black side, whereas for another rat, C5 would be rewarded on the white side and C4 on the black. Generalization testing proceeded immediately after a performance criterion (80% correct during 40 randomized trials) was reached over four sequential days. On each test day, rats were first given 20 randomized reminder trials of the task and then infused bilaterally into the AONs with either muscimol or saline vehicle. After a 30 minute resting period to allow the drug to diffuse throughout the AONs, rats were placed back into the waiting area and then given 3 test trials in only one context per day. During these test trials, one dish contained unscented bedding, one dish contained bedding scented with any one of the odors in the series C2 – C7 (the *test odor*), and neither dish was rewarded. The order of test odor presentations was randomized across rats, in such a manner that each rat was tested on all odors in the series. Rats were given 30 seconds find the reward. The time spent digging in the test dish was recorded using a stopwatch.

#### Experiment 3: Roles of AON and vHC in contextually-cued odor and texture discrimination

To measure the degree to which projections from the vHC and AON are differentially involved in contextually-cued conditional discrimination tasks based on different sensory modalities, a separate cohort of four rats was trained to perform odor- and texture-based conditional discrimination tasks simultaneously. Rats were trained on the contextually-cued odor task using cumin and oregano with context-odor associations counterbalanced between rats. Once rats reached criterion on the conditional odor discrimination, they began training on texture stimuli. Rats were trained on the contextually-cued texture discrimination using corn and paper bedding with context-texture associations counterbalanced between rats. After reaching criterion on the conditional texture discrimination, they then were trained to perform odor and texture tasks simultaneously. On average, a rat would perform 10 texture and 10 odor trials on each side of the box, with all trials randomized (40 trials a day). After rats had maintained criterion performance for four subsequent days, they were tested after being infused (1) with saline in both the AON and vHC, (2) with saline in AON and muscimol in vHC, or (3) muscimol in AON and saline in vHC. All rats were tested under all three drug conditions; the order of drug infusions was randomized among rats.

#### Data Analysis

Performance in the contextually cued odor discrimination, non-contextually cued odor discrimination, and contextually cued texture discrimination tasks (Experiments 1 and 3) was analyzed by comparing the proportions of correct trials for each experiment between saline- and muscimol-infused rats using one-way independent measures ANOVA.

For non-contextually cued generalization results (Experiment 2a), we analyzed digging times during unrewarded trials using a mixed model ANOVA, with *odorant* (C3-C8) as a within-subjects factor and *drug* (saline, muscimol) as a between-subjects factor. The alpha criterion was set to 0.05.

Contextually-cued generalization data (Experiment 2b) were analyzed using a mixed model ANOVA of digging times, with *odorant* (C2-C7) as a within-subjects factor and *drug* condition as a between-subjects factor. In post hoc testing, we asked subsequently whether a significant effect of odor was observed under each drug condition (one-way repeated-measures ANOVA, Wilks’ lambda criterion, Bonferroni-corrected alpha = 0.025 to compensate for multiple comparisons).

#### Computational modeling

The computational model comprised single-compartment probabilistic leaky integrate-and-fire neurons, with the exception of mitral cells (MCs) which were modeled with two compartments. The equations defining these neurons were adapted from previous models (de Almeida et al 2013, Devore et al 2014, Linster et al 2007, Linster et al 2009, Mandairon et al 2014). Model parameters are listed in Table 2. Changes in membrane voltage v(t) over time in each compartment are described by equation 1:

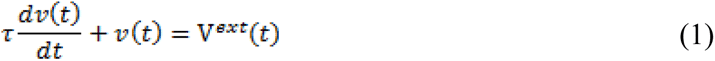

**Table 2:** Computational modeling parameters. Membrane time constant: τ; resting membrane potential: V_rest_; spiking threshold: θ^min^; saturation threshold: θ^max^; synaptic weight: w; reversal potential : E_N_; rise time : τ_1_; decay time : τ_2_.

|  |  |
| --- | --- |
| Olfactory Sensory Neuron (OSN) | $\tau = 1\text{ms}$; $V_{rest} = -65 \text{ mV}$; $\theta^{min} = -65\text{mV}$; $\theta^{max} = -55\text{mV}$. |
| Mitral (MC) | $\tau = 4\text{ms}$; $V_{rest} = -65 \text{ mV}$; $\theta^{min} = -64\text{mV}$; $\theta^{max} = -55\text{mV}$ |
| Periglomerular (PG) | $\tau = 2\text{ms}$; $V_{rest} = -65 \text{ mV}$; $\theta^{min} = -65\text{mV}$; $\theta^{max} = -60\text{mV}$ |
| Granule (GC) | $\tau = 4\text{ms}$; $V_{rest} = -65 \text{ mV}$; $\theta^{min} = -66\text{mV}$; $\theta^{max} = -60\text{mV}$ |
| External tufted (ET) | $\tau = 8\text{ms}$; $V_{rest} = -65 \text{ mV}$; $\theta^{min} = -64\text{mV}$; $\theta^{max} = -60\text{mV}$. |
| Pyramidal (Pyr) | $\tau = 10\text{ms}$; $V_{rest} = -65 \text{ mV}$; $\theta^{min} = -68\text{mV}$; $\theta^{max} = -55\text{mV}$ |
| OSN to PG | $w = 0.0015$; $E_N = +70\text{mV}$; $\tau_1 = 1\text{ms}$; $\tau_2 = 2\text{ms}$ |
| OSN to MC (apical) | $w = 0.015$; $E_N = +70\text{mV}$; $\tau_1 = 1\text{ms}$; $\tau_2 = 2\text{ms}$ |
| OSN to ET(apical) | $w = 0.0015$; $E_N = +70\text{mV}$; $\tau_1 = 1\text{ms}$; $\tau_2 = 2\text{ms}$ |
| PG to MC (apical) | $w = 0.002$; $E_N = -5\text{mV}$; $\tau_1 = 2\text{ms}$; $\tau_2 = 4\text{ms}$ |
| ET to MC (apical) | $w = 0.015$; $E_N = 70\text{mV}$; $\tau_1 = 1\text{ms}$; $\tau_2 = 2\text{ms}$ |
| MC (soma) to GC | $w = 0.001$; $E_N = +70\text{mV}$; $\tau_1 = 1\text{ms}$; $\tau_2 = 2\text{ms}$; $p = 0.2$ |
| GC to MC (soma) | $w = 0.025$; $E_N = -10\text{mV}$; $\tau_1 = 2\text{ms}$; $\tau_2 = 4\text{ms}$; local only |
| Pyr to ET | $w = 0.02$; $E_N = +70\text{mV}$; $\tau_1 = 1\text{ms}$; $\tau_2 = 2\text{ms}$; $p = 0.2$ |
| Pyr to GC | $w = 0.02$; $E_N = +70\text{mV}$; $\tau_1 = 1\text{ms}$; $\tau_2 = 2\text{ms}$; $p = 0.2$ |

where τ is the membrane time constant and *V^ext^*(*t*) is the voltage change resulting from external input over time.

Each one of the voltage changes due to external inputs *V^ext^* is a result of the synaptic weight of the connection from neuron *j* to neuron *i* (*w_ij_*) and the respective synaptic conductance in cell *i* at time *t*, *g_i_*(*t*). *E_N,ij_* is the Nernst potential of the synaptic current, and *v_i_*(*t*) is the membrane potential of the postsynaptic neuron *i*, as described in equation 2:

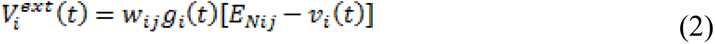

The communication between neurons is mediated by discrete spikes. The spiking output *F(v)* of a given neuron *i* is a function of its membrane potential *v* and the minimal threshold *θ^min^* and saturation threshold *θ^max^* of the output function, where *F_i_*(*v*) = 0 if *v*≤*θ^min^*, *F_i_*(*v*) = 1 if *v*≥*θ^max^*, and *F_i_(v)* increases linearly between θ_min_ and θ_max._

The time course of the conductance change *g_i_(t)* is calculated as:

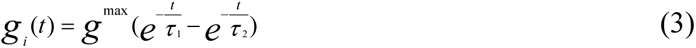

where *g^max^* is a constant representing the maximum conductance of a given channel and set to 1 (synaptic strengths are instead scaled by the synaptic weight *w*), while τ_1_ and τ_2_ are the rising and falling time constants of this conductance. After firing, the voltage of each neuron is reset to V_rest_.

Computational modeling followed the principles outlined in Mandairon et al. (2014). Simulated classes of neurons included olfactory sensory neurons (OSN), external tufted cells (ET), periglomerular cells (PG), mitral cells (MC), granule cells (GC) and AON pyramidal cells (Pyr). In the model, 100 of each cell type are represented. OSNs make excitatory synapses onto ET, PG and MC cells within a given glomerular column. Within these columns, ET cells excite MCs, and PG cells inhibit MCs. MCs excite a subset of randomly chosen GCs (20% connectivity) while GCs inhibit MCs within the same column, reflecting the fact that inhibition is most effective when delivered proximal to the MC soma (David et al 2008, McIntyre & Cleland 2016). Simulated odorants activated a randomly selected subset of 10% of the OSNs using a normal distribution of affinities; this incoming odor representation then was further modulated by interactions with local interneurons (PG, ET and GC cells).

Pyramidal cells in the simulated AON represent contextual information conveyed by the vHC (not modeled), wherein each simulated spatial context (*black* or *white*) activates a distinct subset of 10% of AON Pyr neurons. Tasks in which context was irrelevant were represented by spontaneous activity in AON Pyr neurons, whereas muscimol lesions of AON were represented by no output from AON Pyr neurons. These Pyr neurons synapse onto both GCs and ETs with excitatory synapses, as demonstrated experimentally (Boyd et al 2015, Markopoulos et al 2012), with a 20% probability using a uniform distribution. As a consequence, contextual information modulates odor representations by introducing both an excitatory (ET) and an inhibitory (GC) bias onto MC activity.

## Code accessibility

The code/software described in the paper will be made freely available online at http://modeldb.yale.edu/.

## Results

We first show that a functioning AON is necessary for rats to perform a well-learned contextually-cued odor discrimination task, but not for a simple odor discrimination in which context is not relevant. We then show that this function of the AON is specific to odor-based but not texture-based contextually-cued discrimination (Experiment 1). We then test to what extent context can modulate odor perception using a well-defined olfactory generalization task. We show that the AON does not modulate odor perception unless a contextual cue is necessary to solve the task (Experiment 2). Computational modeling demonstrates how AON responses to behaviorally relevant contexts can modulate OB odor responses such that MC population responses to a given odor are quite different in different contexts. Finally, we show that information from vHC is important for both odor- and texture-based conditional discrimination, whereas AON lesions impair only odor-related tasks (Experiment 3).

### Experiment 1: A functioning AON is required for contextually-cued conditional odor discrimination

To examine to what extent the AON is involved in contextually-cued conditional odor discrimination, we tested rats on their ability to perform a well learned contextually-cued conditional odor discrimination task with and without a functioning AON (Experiment 1a). The visuospatial contexts comprised a three-chamber test arena with one *black* and one *white* context chamber surrounding an intertrial chamber in the middle (Figure 1Ai). Rats first were trained to retrieve a sugar pellet from a dish scented with odor A but not odor B in one of these visuospatial contexts and the reverse in the other context (see *Methods*). After reaching criterion on the task, rats were bilaterally infused with muscimol or saline vehicle into the AONs and then tested on their task performance. Rats performed significantly less well on the contextually-cued task when muscimol was infused into the AON (effect of *drug* on the proportion of correct trials: F(1, 6) = 38.124, *p* = 0.003; Figure 1B, *contextual odor*), showing that AON function is necessary for performance in this task. When rats were tested in a task not involving contextual cues (Experiment 1b), using a box with a single test chamber (Figure 1Aii), muscimol inactivation of the AONs did not affect performance (effect of *drug*: F(1,4) = 0.10, *p* = 0.786; Figure 1B, *non-contextual odor*). To test whether AON involvement in this task was specific to olfactory stimuli, rats then were trained to perform an identical contextually-cued discrimination task using the texture of the digging medium (corn bedding versus paper bedding) rather than odorant as a cue to which dish contained the sugar reward (Experiment 1c). After reaching criterion on this task, rats were bilaterally infused with either muscimol or saline into the AONs and then tested. Each rat was tested twice with each drug. In contrast to their strong effect on odor-dependent contextual discrimination, no effect of muscimol infusions into AON was observed in the texture-dependent discrimination task (F(1, 14) = 1.912, *p* = 0.188; Figure 1C, *contextual texture*). The location of infusion cannulae was verified histologically after the end of behavioral experiments (Figure 1C).

### Experiment 2: The AON modulates odor perception only when contextual information is task-relevant (*Figure 2*)

To assess whether and when AON regulates odor perception, rats were trained to discriminate between pentanal (C5) and hexanal (C6) in a non-contextual task (Figure 1Aii) and subsequently tested on how they generalized between the trained odors and novel odors of varying similarity (Experiment 2a). Straight-chain aliphatic molecules with increasing carbon chain lengths are treated as sequentially similar in behavioral tests, such that rats trained on a five-carbon aldehyde (C5) will generalize more strongly to a four-carbon or six-carbon aldehyde (C6, C4) than to a three- or seven-carbon aldehyde (C3, C7) (Cleland et al 2002), although the extent of generalization depends substantially on learning (Cleland et al 2011, Cleland et al 2009). On a given test day, rats were first trained to discriminate odors C5 and C6 over 20 trials, with half of the rats being rewarded for C5 (C5+/C6-) and half for C6 (C6+/C5-). After the 20 trials, rats were bilaterally infused with muscimol or saline vehicle into the AON and then tested in non-rewarded trials with straight-chain aldehyde odorants ranging from 3 (C3) to 8 (C8) carbons in the aliphatic chain. The time spent digging in these unrewarded trials was recorded as a measure of generalization from the trained odors. Figure 2A illustrates the pattern of perceptual generalization expected from discrimination training: rats dig in response to odors similar to the rewarded odor and do not dig in odors more similar to the non-rewarded odor, with the different contingencies of the two trained odorants forming a sharp cutoff. Rats with muscimol infusions into their AONs showed the same generalization pattern in this task as vehicle-infused control animals, demonstrating that impairing AON function has no effect on odor generalization per se (Figure 2A). Analyzing digging times with *odorant* as a within-subjects factor and *drug* as a between-subjects factor, we found a significant effect of *odorant* (F(5, 4) = 26.025, *p* = 0.004) but no interaction of *odorant* × *drug* (F(5,4) = 1.454, *p* = 0.369), indicating that muscimol infusions into the AON did not affect odor generalization in this task. These results show that odor perception per se is not modulated by AON activity.

**Figure 2.**
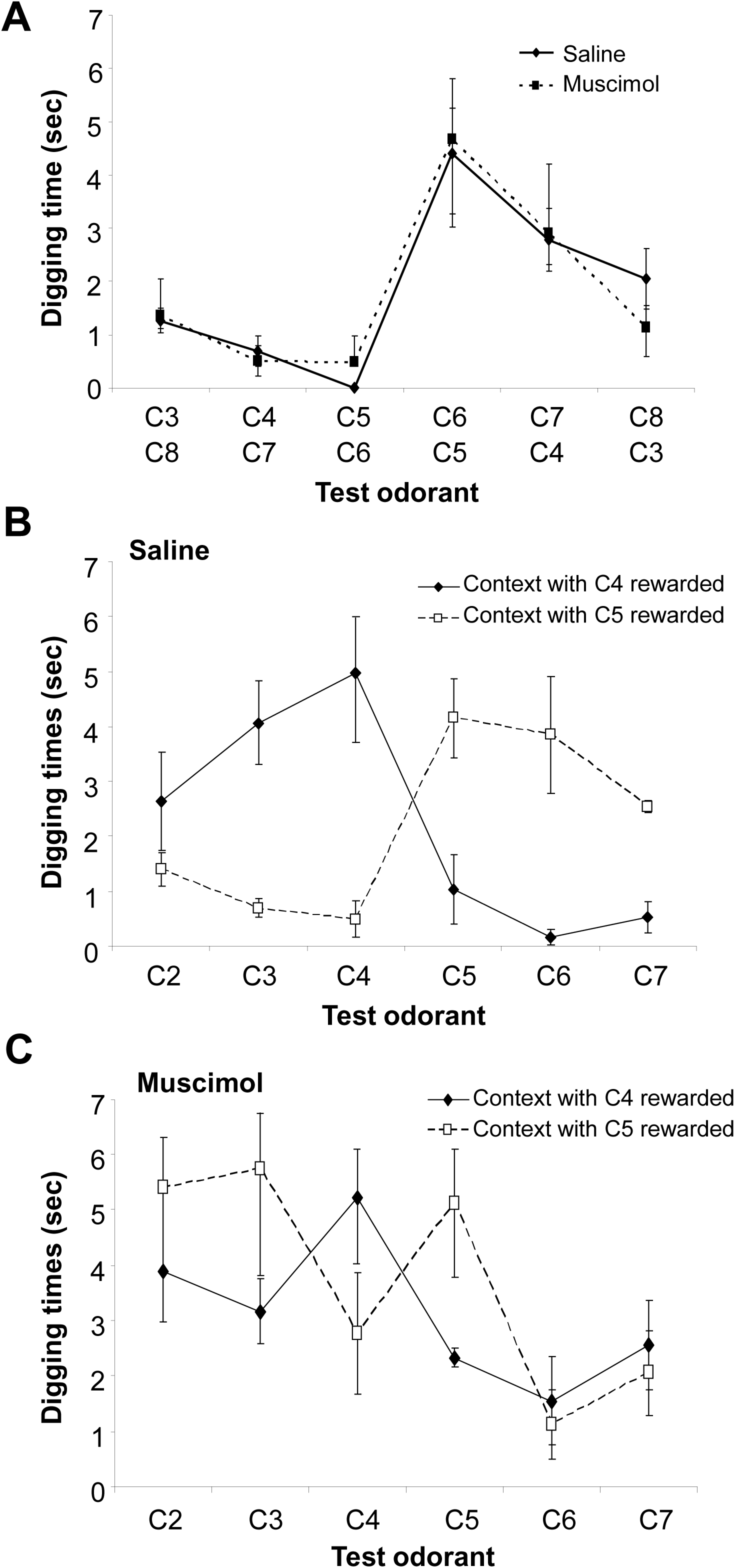
Role of AON in odor perception. The graphs show mean (+/-standard error) digging times during unrewarded test trials. **A**. Digging times during unrewarded test trials in rats trained in a non-contextually cued odor discrimination task after saline or muscimol infusions. All rats dug most in the rewarded odorant and least in the non-rewarded odor, and exhibited expected generalization gradients to novel odorants with varying degrees of perceptual similarity. **B**. Perceptual generalization in both contexts in saline-infused rats. The graph shows digging times as a function of unrewarded test odorant when rats were tested in the context in which they were rewarded for C4 or the one in which they were rewarded for C5. The degrees of perceptual generalization are similar in each context and expressed as digging for odorants similar to the rewarded odor and as the absence of digging in odorants similar to the nonrewarded odor. **C.** Perceptual generalization in both contexts in muscimol-infused rats. The graph shows digging times as a function of unrewarded test odor when rats were tested in the context in which they were rewarded for C4 or the one in which they were rewarded for C5. In this case rats treated all odors similarly, with no clear generalization from the rewarded or unrewarded odors.

Using novel odorants (aliphatic butyrates), we then tested to what degree task-relevant contextual information modulates odor perception following contextually-cued training (Experiment 2b). For example, a rat was conditioned to expect a reward in butyl butyrate but not pentyl butyrate (C4+ / C5-) on the *white* side of the box, and the reverse (C5+ / C4-) on the *black* side of the box. After learning this task to criterion, the rats’ generalization to novel odors was tested on each side of the box, with the expectation that context would modulate odor perception and alter generalization gradients. After presenting 20 reminder trials with randomized trials in each context, rats were bilaterally infused in the AONs with saline vehicle, and then performed three unrewarded trials a day in a given context with a series of straight-chain aliphatic butyrates ranging from two to seven carbons in the aliphatic chain. As predicted, the rats responded differently to novel odorants depending on the visuospatial context, as if that context cued a clear memory of which odorant was rewarded (Figure 2B); specifically, individual rats responded differently in the *black* and *white* contexts, according to which odor had been rewarded for them in that context. However, after muscimol infusions into the AONs, rats responded statistically similarly to all odorants in both contexts, suggesting that, in the absence of this contextual information from AON, generalization between odors was disrupted, presumably because no clear predictor of reward could be established (Figure 2B). A repeated measures ANOVA with *digging time* as the dependent variable, *drug* condition as the between-subjects factor and test *odorant* as the within-subjects factor revealed a significant effect of *odorant* (F(5, 16) = 4.085; *p* = 0.014) as well as a significant interaction between *odorant* × *drug* (F(5, 16) = 2.960; *p* = 0.044). These results show that relative digging times in the test odors are dependent on drug treatment. Further post hoc tests revealed a significant effect of *odorant* on digging times in the saline (F(5,6) = 5.252; *p* = 0.034) but not muscimol condition (F(5,6) = 0.696; *p* = 0.646), showing that muscimol-treated animals did not significantly differ in their responses to the test odorants whereas saline-treated animals did. For saline-infused animals, these results present a classical generalization pattern (Figure 2B). In contrast, when AON function was impaired via muscimol infusions, rats retained their capacity to differentiate between the rewarded and unrewarded odorants, but otherwise generalized to all test odorants indiscriminately. These results show that visuospatial context can modulate odor perception if it is task-relevant, and that a functioning AON is required to convey this contextual information.

### Computational modeling illustrates how the AON-dependent contextual modulation of odor representations can underlie differential generalization gradients

We then simulated contextually cued and non-contextually cued processing of odors using our established olfactory bulb model to which contextual information in the AON had been added. With respect to contextual information, three scenarios were modeled: (1) contextual information is not required for task performance (*irrelevant context*); in this case AON activity consisted of spontaneous activity alone; (2) context is task-relevant; in this case contextual representations activated a consistent, non-overlapping subset (10%) of AON pyramidal cells for each of two contexts (*black context* or *white context*); and (3) AON was lesioned by muscimol (*no context*); in this case AON cells were silent. Series of sequentially similar odors as used in behavioral generalization experiments (Cleland et al 2002, Cleland et al 2009, Mandairon et al 2006a, Mandairon et al 2006b) were simulated by using normal distributions of sensory neuron activations correlated by 0.85 for each neighboring odor in the sequence. Generalization gradients in the model were calculated as pairwise correlations between average MC responses to simulated odorants (Linster & Cleland 2002).

Figure 3 shows examples of simulated spike trains and membrane voltages for neurons in the model in response to presentation of an odorant stimulus (*blue bar*) with irrelevant context (first half of time shown) and in response to odorant presentation within a relevant context (*red bar*; second half of time shown). Sensory neurons (OSNs) respond to odor stimulation with various mean spike rates modulated by respiration and convey this information to ET, PG and MC cells. Pyramidal cells in the AON fire spontaneously in the absence of task-relevant context and a subset are activated strongly by a given context. This contextual activation converges onto ET and GC neurons along with odorant-related afferent activation; consequently, both ET and GC cells exhibit consistently modulated odor responses when contextual information is present (Figure 3, ET, GC, *blue arrows*). Their respective excitatory and inhibitory effect on MC principal neurons enables specific contexts to both generate or heighten some MC responses (Figure 3, MC, *blue arrows*) while inhibiting others (Figure 3, MC, *red arrows*), thereby influencing the odorant representations exported from the OB to other regions of the brain.

**Figure 3.**
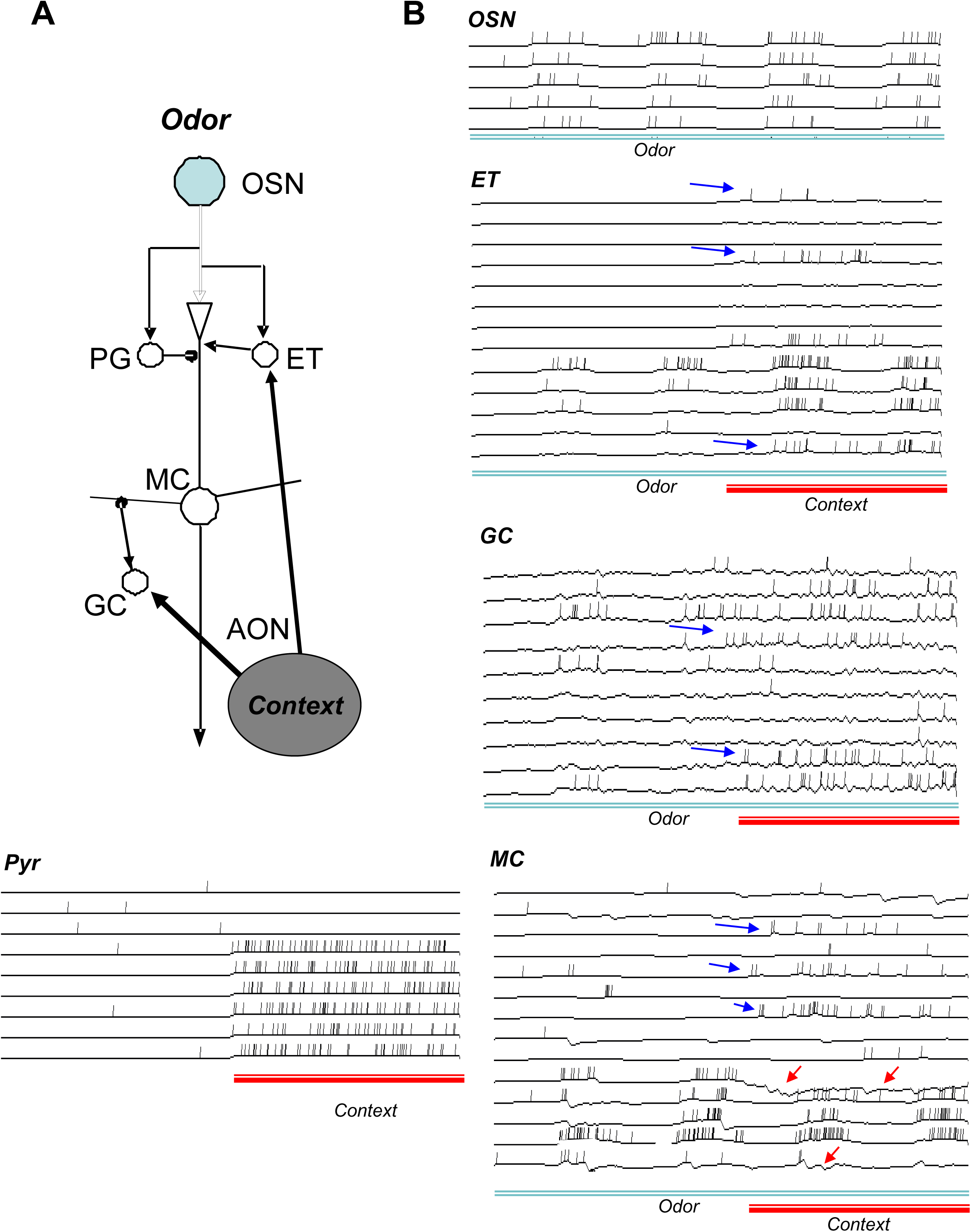
Model schematic and neuronal responses. **A**. Model schematic: The model is organized into 100 columns with one sensory (OSN), periglomerular (PG), external tufted (ET), mitral (MC), and granule (GC) cell associated with each column. Connections between glomerular interneurons and mitral cell dendrites are within a column, connections from mitral to granule cells are distributed equally across columns with a 20% probability, and connections from granule to mitral cells are local within each column to reflect the local efficacy of inhibition. Feedback inputs from AON pyramidal cells are distributed randomly to ET and GC cells with a uniform 20% probability. **B.** Examples of model neuronal responses to an odor (*blue bar*, 800 ms) and an odor+context (*red bar*, 400 ms) are shown for each cell type in the model. Blue arrows designate examples of increased activity in the presence of context and odor as compared to odor alone, red arrows designate examples in which the addition of context reduces or inhibits odor responses.

We then measured the extent to which contextual feedback from the AON can modulate odor representations in the OB (Figure 4). We randomly selected one odor to stimulate OSNs, and ran the model either with spontaneous activity in the AON (*irrelevant context*) or with one of two non-overlapping sets of contextual information (*white context, black context*) activating a corresponding 10% of AON neurons above spontaneous activity). The *irrelevant context* condition in these simulations corresponds to rats learning the non-contextually cued odor discrimination task (Experiment 1b), and the two contexts correspond to learning a contextually cued odor discrimination task in which context matters (Experiment 1a). To ensure that the results were not dependent on a specific network architecture, we ran 10 instances of the network with 50 repetitions each. Each odor representation at the output of the OB was represented as a 100-dimensional vector of mean MC responses to that odor during stimulation; pairwise comparisons between representations were computed as correlations between their activity vectors. Whereas the overlap between successive simulations of a given odor under *irrelevant context* conditions is very high (Figure 4A, *irrelevant context*) the overlap between representations of the same odor under different contextual circumstances is significantly lower (*irrelevant context / white context*) and the overlap between representations of the same odor biased by two separate contexts is less than 40% (*white context / black context*). Comparing these correlations showed a significant effect of contextual condition with F(2, 104) = 316.167; *p* < 0.001) and significant pairwise differences between all conditions (LSD, *p* < 0.001 in all cases). These results show that the receipt of relevant contextual information from the AON, using a random distribution of AON projections to inhibitory GCs and excitatory ET cells, can significantly bias MC activation patterns such that odors learned in each context could create quite distinct percepts. Accordingly, if an odor-reward association is learned in a given salient context, it may not necessarily be fully recalled outside of that context.

**Figure 4.**
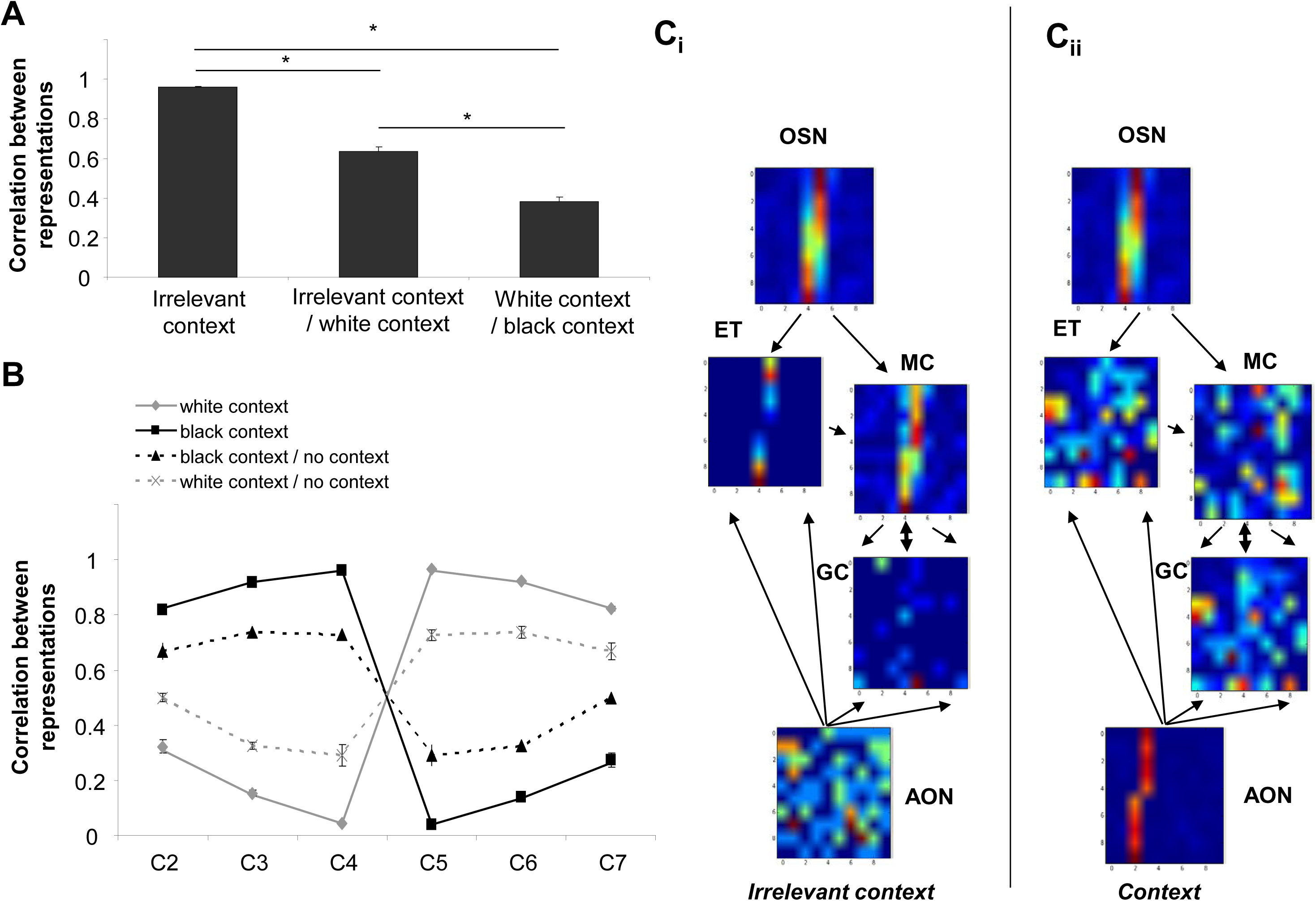
Computational modeling results. **A**. Pairwise correlations between MC population responses to the same odor presented under different conditions: *irrelevant* context, *white* context, or *black* context. Correlations between representations simulated under the same condition (*irrelevant* context) are very high. Correlations between representations simulated under two different conditions (*irrelevant context/white context* and *white context/black context*) are considerably lower. These simulations show that contextual information can bias OB representations. *denote significant differences between conditions. **B.** The graph shows correlations between odor responses simulating the behavioral generalization experiments. Odor C4 has negative valence in the *white* context, and odor C5 has positive valence (valences are reversed in the *black* context). Correlations between odors C4/C5 and similar odors follow expected generalization gradients when these odorants are presented under the same conditions in which they were learned (*white* or *black* context). In contrast, when odors C2-C7 are simulated under AON lesion (*no context*) and correlated with C4 or C5 under white or black context, the generalization gradients become flat and differences between negative valence odors and positive valence odors are less pronounced. **C.** Flow of processing in the model. Average spiking activity of each class of neurons are depicted as 10×10 heatmaps with warmer colors signaling higher activity. In the no context information (panel Ci), a randomly chosen odorant activates a subset of OSNs and this activity is transmitted to ET and MC cells. MC cells connect randomly to GC cells and generate distributed activity among them. In the no-context situation, AON cells are weakly activated in a noisy and random fashion. In the presence of contextual information (panel Cii), a subset of AON cells is strongly activated, thereby substantially biasing the activation of GC and ET cells and consequently transforming MC cell representations to include contextual information.

These simulations suggest that contextual information from the AON can bias OB representations to generate substantially different activation patterns in response to the same odorant under different contextual circumstances. Alternatively, the OB can be thought to construct multimodal representations comprising both odor and (visuospatial) contextual information. To assess whether AON contextual inputs can bias odor representations in the model sufficiently to explain the results of Experiment 2, we used the model to assess the effect on generalization gradients. Behavioral experiments were modeled by computing the OB output vector in response to a rewarded odor (C4 in black context and C5 in white context) in the presence of contextual information conveyed by the AON. We then calculated pairwise correlations between the rewarded odor in its original context and all odorants (C2–C7) in either the same context or no context. To reflect the fact that in context white C5 was rewarded but not C4, the difference between maximal correlation coefficient (1.0) and the calculate correlation coefficient (1.0 – correlation coefficient) was plotted for the non-rewarded odor (C4) and odors closest to it (C3 and C2). Similarly, in black context, 1-correlation coefficient was plotted for the non-rewarded odor (C5) and odors closest to it (C6 and C7). This resulted in the rewarded odor being highly correlated with itself when presented in the same context, but not when presented in the absence of context. Similarly, the correlation between the non-rewarded odor and the rewarded odor in the same context was very low (close to zero, reflecting a very low behavioral response), but higher when compared to the no-context simulation. Generalization among odorants was very similar when tested under the same circumstances (Figure 4B; white context and black context). However, if we compared representations learned in a salient context to those obtained in a no-context simulation (mimicking AON lesions), the generalization gradients were flattened, and reminiscent of those obtained following temporary AON lesions (Figure 2). In this situation, the absence of contextual information rendered the odorant-evoked OB representation less similar to that previously obtained in response to that same odorant presented within a salient context. These simulations showed that contextual information can bias or change OB odor representations sufficiently for generalization gradients to be context dependent.

Figure 4C illustrates the mechanisms of the OB model with respect to the contextual modulation of odorant representations. Each heat map is a 10×10 depiction of the mean firing rates of a population of model cells in response to one odor in the *irrelevant context* condition (Ci) and in a *context* condition (Cii), with warmer colors depicting higher spike rates. When no contextual information is present (*irrelevant context*), the AON exhibits spontaneous background activity. OSN activation patterns are clearly reflected in ET and MC activity, whereas GCs are activated primarily by MC activity. When a subset of AON pyramidal neurons is strongly activated by a salient context (*context*), ET, GC, and MC neurons are driven (directly or indirectly) by both odorant and contextual inputs; consequently, ET and MC activity patterns no longer clearly reflect OSN activation. In essence, the OB output representation now embeds both olfactory and contextual information. Importantly, however, this embedding does not comprise a simple addition of those signals, as AON activates both inhibitory and excitatory interneurons, leading to mixed effects on MC representations.

### Experiment 3: The vHC is required for contextually cued discrimination irrespective of stimulus modality, whereas AON function is specific to olfactory stimuli

We hypothesized initially that projections from vHC to AON convey contextual information needed to cue the animal to which odor is rewarded in a conditional discrimination task. To test the extent to which each of these structures is domain general for context or modality-specific for olfaction, a new cohort of rats with bilateral cannulae placed into the AON and vHC was trained to simultaneously perform contextually-cued odor and texture discriminations. Rats learned to associate a reward with a particular odor or texture within a given context. After reaching criterion, rats underwent 40 trials in which the visuospatial context (*black* or *white*) and the stimulus modality (*odor* or *texture*) were randomly combined. On test days, rats were infused with muscimol or saline vehicle in the AON or ventral hippocampus, yielding three *drug* conditions: AON_muscimol_/vHC_vehicle_, AON_vehicle_/vHC_muscimol_, and AON_vehicle_/vHC_vehicle_ controls. Individual rats were run under each of these drug conditions in an order that was randomized among rats. The mean percentage of correct trials during testing on this task under the different drug conditions is depicted in Figure 5A. When infused with saline vehicle in both the AON and vHC, rats performed at above 80% correct on both odorant and texture stimuli. Rats infused with muscimol in the AON and saline in the vHC were impaired in the odorant but not the texture discrimination task, replicating the findings of Experiment 1 (Figure 1B). In contrast, rats infused with muscimol in vHC (and saline vehicle in the AON) were impaired in both odorant and texture discriminations. An ANOVA with *drug* condition and stimulus *modality* as between-subjects effects revealed significant effects of both *drug* (F(2, 30) = 17.59, *p* < 0.001) and *modality* (odor or texture: F(1, 30) = 13.24; *p* = 0.001) as well as a significant interaction between these terms (F(2, 30) = 25.05; *p* < 0.001). These results indicate that the three drug infusion conditions affected performance on odor and texture trials differently. Using post hoc pairwise comparisons (LSD, alpha = 0.016 to compensate for three comparisons), we then compared the rats’ odor and texture discrimination performance under each drug infusion condition. Rats infused with saline in both structures performed equally well on each set of trials (Figure 5A, *left panel*; F(1 10) = 2.083; p = 0.498), whereas rats infused with muscimol in the AON and saline in the vHC performed significantly less well for odor stimuli compared to texture stimuli (Figure 5A, *center panel*; F(1, 10) = 121.622; p < 0.001). Rats infused with muscimol in the vHC and saline in the AON performed equally badly for both odor and texture stimuli (Figure 5A, *right panel*; F(1, 10) = 0.522; p = 0.486). These results demonstrate that a functioning vHC is required to convey contextual information in both olfactory and tactile contextually-cued discrimination tasks, whereas a functioning AON is only required for the olfactory version of this task.

**Figure 5.**
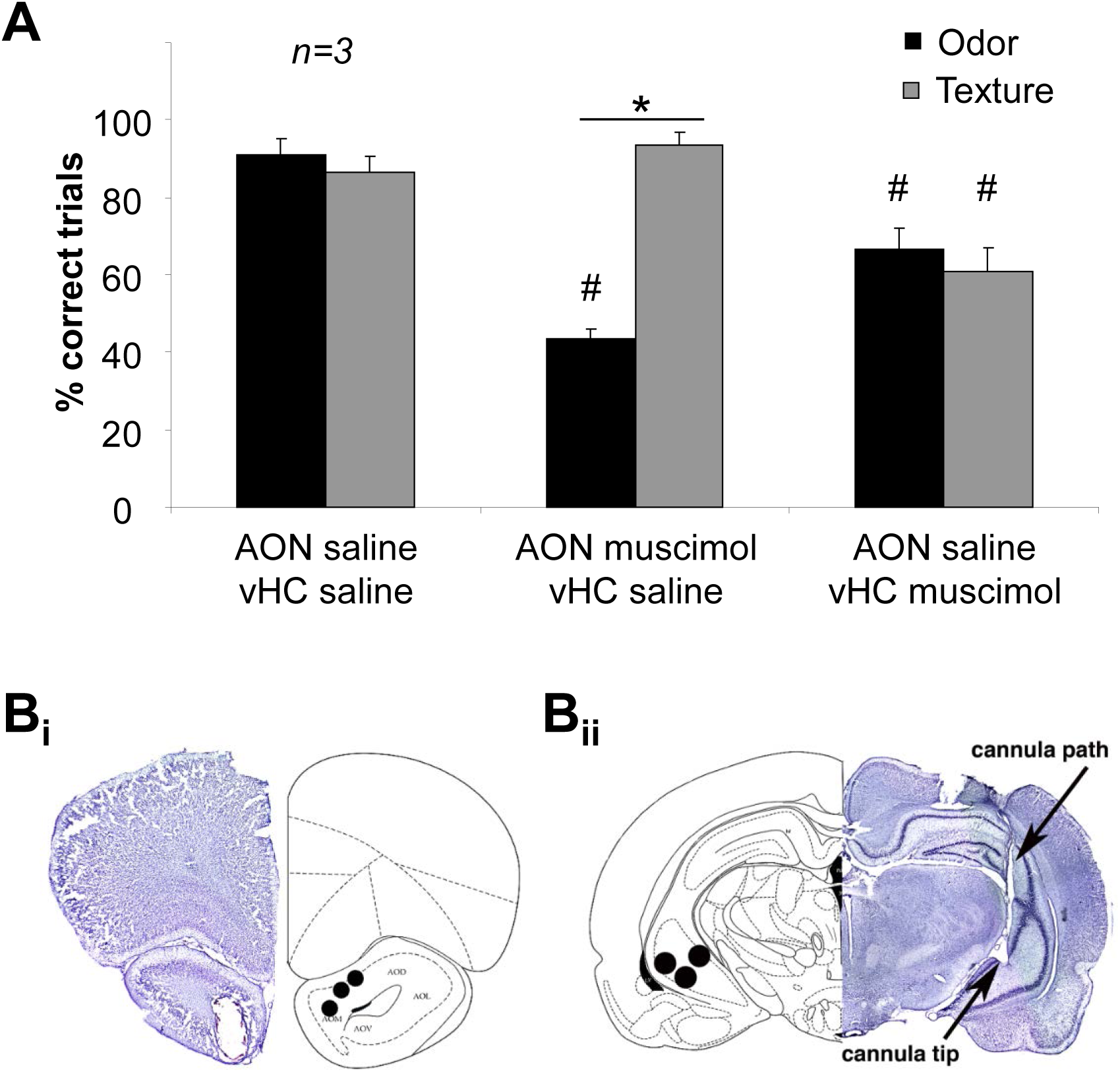
Roles of AON and vHC in contextually-cued odor and texture discrimination. **A.** The graph shows the mean percentage of correct responses to odor and texture trials as a function of drug infusions. * indicates a significant difference between texture and odor in the same drug condition and # indicates a significant difference from the vehicle-only control condition. **B.** Histological verification of cannula placements in the AON (Bi) and vHC (Bii).

## Discussion

We here describe a novel function of the anterior olfactory nucleus in odor learning and olfactory processing. Specifically, we show that environmental context can exert a profound influence on olfactory perception, resulting in context-specific generalization gradients among odorants. Contextual modulation was abolished by inactivation of the AON, indicating that the AON is a critical conduit of information about the visuospatial context – a role that is clearly consonant with the AON’s anatomical connections to olfactory and hippocampal structures (Aqrabawi & Kim 2018b, Brunjes et al 2005, Haberly & Price 1978). In particular, the AON receives a well-defined projection from the ventral hippocampus (Aqrabawi et al 2016, Aqrabawi & Kim 2018b, van Groen & Wyss 1990). The hippocampus is known to be involved in processing contextual information (reviewed by Smith and Bulkin, (2014)) and the ventral hippocampus specifically has been implicated in the contextually-cued conditional discrimination paradigm used herein (Komorowski et al 2013). Once rats have learned this task, temporary lesions of the AON impair performance if, and only if, visuospatial contextual cues are essential to olfactory task performance, suggesting that contextual information is not available to the rats when identifying the rewarded odor. Interestingly, when rats perform the same contextually-cued task using a texture rather than an odorant stimulus to find the reward, lesions of the AON do not impair performance whereas vHC lesions do. This suggests a framework in which contextual information represented in the vHC is specifically conveyed to the AON when required for performance of an olfaction-dependent task.

We further show that rats can express multiple context-specific perceptual generalization profiles to the same odorant. Previous experiments from our lab and others led to the hypothesis that generalization among odorants is determined by neural circuitry as early as the olfactory bulb, and is governed by physical similarities among odor representations conveyed to the OB by olfactory sensory neurons (Cleland et al 2002, Cleland et al 2009, Linster & Hasselmo 1999, Linster et al 2001, Mandairon & Linster 2009, McNamara et al 2008). Perceptual generalization is easily modulated by pharmacological manipulations of OB function as well as through learning and experience (Cleland et al 2009, Devore & Linster 2012, Mandairon & Linster 2009). We therefore hypothesize, as depicted in the computational model, that contextual cues directly modify odorant representations in the OB. That is, projections from the AON to the OB would serve to bias odor representations in the OB according to salient contextual factors, altering odor representations and their similarities as a function of non-olfactory context. However, when contextual cues are not task relevant during learning, AON inputs become irrelevant and do not strongly influence odorant representations in OB; we modeled this as weak and random activity in AON pyramidal neurons that affected the OB in an unbiased manner and hence would not accumulate over the course of learning. To test these ideas, we used our established model of OB function and generalization among representations (de Almeida et al 2013, Devore et al 2014, Linster & Cleland 2002, Linster et al 2011), adding AON projections to the OB in a manner consistent with experimental findings (Boyd et al 2015, Markopoulos et al 2012) and hypothesizing that task-relevant contextual cues are embedded in the activity profiles of AON pyramidal cells (Mandairon et al 2014), with different visuospatial contexts activating different characteristic ensembles of AON neurons. (In the present version of this model, AON projections to the OB are sparse and randomly distributed, and not plastic). Critically, this implies that activity representations in OB mitral cells – which receive direct input from primary olfactory sensory neurons – are actually fundamentally multimodal, incorporating both olfactory and non-olfactory information. This hypothesis is supported by electrophysiological recordings showing non-olfactory task-related activity in MCs as well as context-dependent changes in MC odor responsiveness (Doucette & Restrepo 2008, Kay & Laurent 1999, Li et al 2017).

In summary, we propose that the AON conveys information about contextual cues from the ventral hippocampus to olfactory structures, specifically the OB. We suggest that information about behaviorally relevant contextual cues modulates and alters MC representations in the OB, merging contextual with afferent olfactory sensory information. This implies that MCs, which receive direct input from primary sensory neurons and project from the OB to multiple targets across the brain, encode and process multi-modal information to inform decision making.

